# Latent developmental potential to form limb-like skeletal structures in zebrafish

**DOI:** 10.1101/450619

**Authors:** M. Brent Hawkins, Katrin Henke, Matthew P. Harris

**Affiliations:** Department of Genetics, Harvard Medical School, Boston, MA, USA; Department of Orthopedic Research, Boston Children’s Hospital, Boston, MA, USA; Department of Organismic & Evolutionary Biology, Harvard University, Cambridge, MA, USA; Museum of Comparative Zoology, Harvard University, Cambridge, MA, USA

## Abstract

The evolution of fins into limbs was a key transition in vertebrate history. A hallmark of this transition is the addition of multiple long bones to the proximal-distal axis of paired appendages. Whereas limb skeletons are often elaborate and diverse, teleost pectoral fins retain a simple endoskeleton. Fins and limbs share many core developmental processes, but how these programs were reshaped to produce limbs from fins during evolution remains enigmatic. Here we identify zebrafish mutants that form supernumerary long bones along the proximal-distal axis of pectoral fins with limb-like patterning. These new skeletal elements are integrated into the fin, as they are connected to the musculature, form joints, and articulate with neighboring bones. This phenotype is caused by activating mutations in previously unrecognized regulators of appendage development, *vav2* and *waslb*, which we show function in a common pathway. We find that this pathway functions in appendage development across vertebrates, and loss of *Wasl* in developing limbs results in patterning defects identical to those seen in *Hoxall* knockout mice. Concordantly, formation of supernumerary fin long bones requires the function of *hoxall* paralogs, indicating developmental homology with the forearm and the existence of a latent functional Hox code patterning the fin endoskeleton. Our findings reveal an inherent limb-like patterning ability in fins that can be activated by simple genetic perturbation, resulting in the elaboration of the endoskeleton.

## Introduction

Changes in appendage structure underlie key transitions in vertebrate evolution. Tetrapod limbs permit mobility via articulation of multiple endochondral long bones facilitated by specialized synovial joints. Diverse limb structures and functions have evolved in tetrapods through digit reduction, phalangeal addition, transformation of wrist and ankle bones, alteration in element proportion, and outright limb-loss. In contrast to limbs, the skeletal pattern of the teleost pectoral appendage is almost invariant, showing a consistent arrangement across diverse lineages within this group of ∼30,000 species. Teleost pectoral fins are composed of dermal fin rays supported at their base by a diminutive endoskeleton (Fig. 1A). The endoskeleton typically consists of four long bones, called proximal radials, arranged side by side along the anterior-posterior (AP) axis, followed distally by small nodular distal radials *(1, 2)*. No known teleost species exhibits more than a single long bone along the proximal-distal (PD) axis (3). The divergent architectures of teleost fins and tetrapod limbs have been maintained within each clade for over 250 *(4, 5)* and 350 million (6) years, respectively.

**Figure 1.**
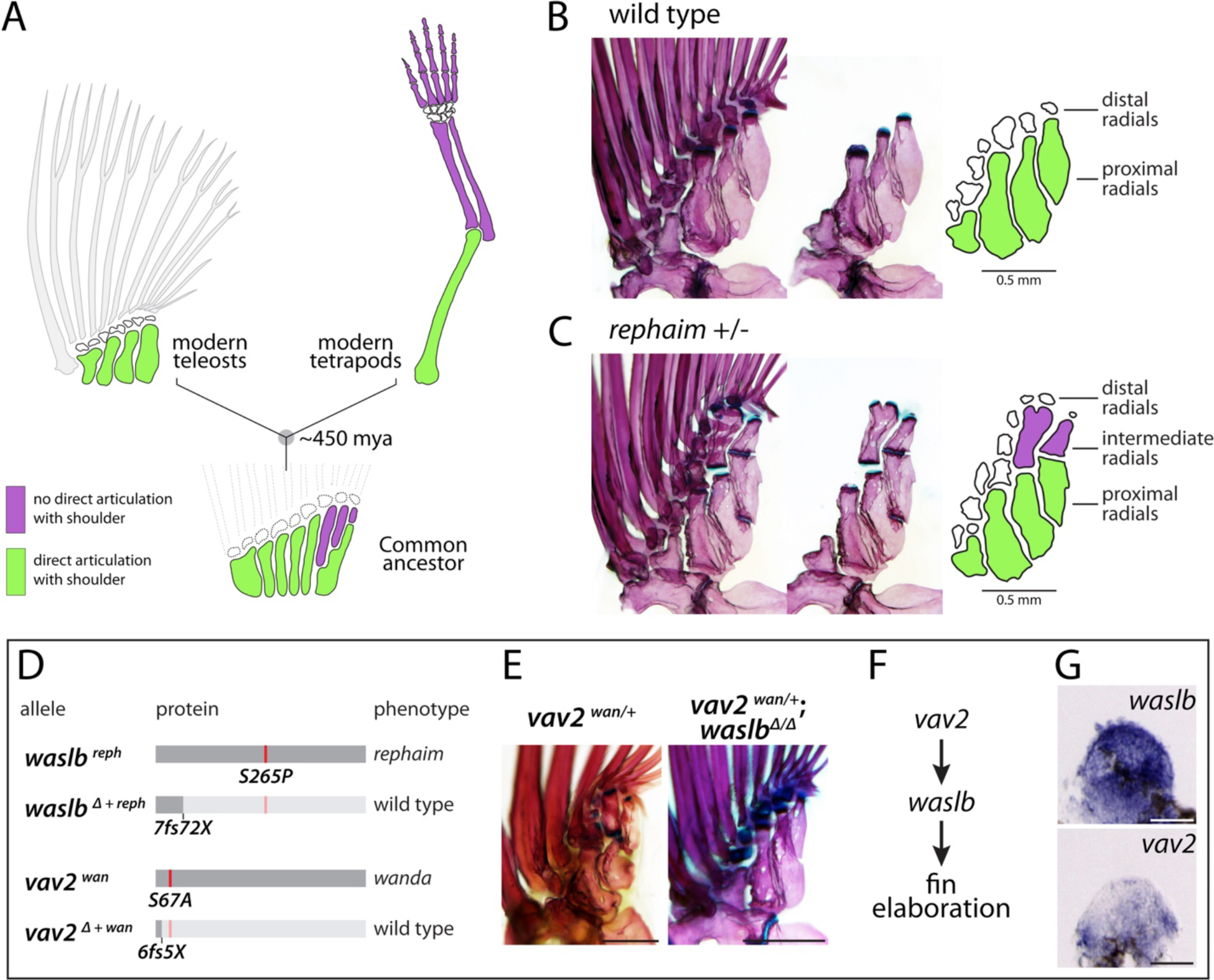
vav2 and waslb gain-of-function mutations reveal the capacity for limb-like development in fins. (**A**) From a common ancestor, teleosts and tetrapods evolved divergent appendage patterns. (**B**, **C**) Adult pectoral fin skeletons of wild-type (**B**) and heterozygous *rephaim* mutants (**C**), shown intact (left), with fin rays and distal radials removed (center), and schematized (right). (**D-G**) Genetic characterization of *reph* and *wan* mutants. (**D**) Schematic of *waslb* and *vav2* alleles found to cause *reph* and *wan* phenotypes, respectively. *Δ + reph* represents allele *mh130*, and *Δ + wan* represents allele *mh138* (Figures S4, S5). (**E**) Epistasis analysis showing suppression of *wan* after the loss of *waslb* function. (**F**) Genetic pathway of vav2, *waslb*, and fin elaboration. (**G**) Expression of *waslb* and *vav2* transcripts in 48 hpf pectoral fin buds. Anterior to left, distal to top; scale bars (B, C, E) 500 μm, (G) 50 μm.

Evidence from fossils suggests that, similar to extant ray-finned fishes, the common ancestor of teleosts and tetrapods had a series of long bones arranged side by side along the AP axis of the pectoral appendage (7). The posterior-most element of this series was segmented along the PD axis, creating end-on-end articulation (Fig. 1A). This structure, called the metapterygium, is not retained in teleost fishes, and is hypothesized to have been elaborated in lobe-finned fishes to form limbs (8) (Figure S1). The progressive acquisition and regionalization of elements along the PD axis of the appendicular skeleton in stem and crown group lobe-finned fishes indicate that the limb is a unique feature in this group not present in ray-finned fishes.

Due to their divergent skeletal patterns, identifying homologous structures between fins and limbs with one-to-one correspondence is difficult. However, despite the morphological disparity between fins and limbs, comparative approaches in developmental genetics have revealed shared pathways required for the normal patterning of these appendages. *Hox* genes, in particular, are key players that specify and reinforce regional identity in appendages (9). Overexpression of *hoxd13a* in embryonic zebrafish causes increased, but unpatterned, chondrogenic tissue formation in the larval pectoral fin *(10)*. Furthermore, deletions of posterior group 13 *HoxA* and *HoxD* genes result in the loss of distal skeletal elements in mouse limbs (wrist/ankle and hand/foot) and zebrafish fins (dermal fin rays), suggesting that these genes provide a regional address for distal elements in the different appendages (11). While *Hox* genes that specify more proximal limb components have been identified in tetrapods, it is unclear if similar mechanisms instruct the patterning of non-distal components of the fin.

Here, through unbiased forward genetic screens in the zebrafish *(12, 13)*, we identify mutants that break the long-standing teleost fin pattern by developing new skeletal elements along the PD axis of the pectoral fin in a limb-like manner. These novel structures are fully differentiated long bones that integrate into the fin musculature and articulate with the fin skeleton. Mapping and epistasis analysis reveal modulation of *vav2* and *waslb* signaling as causing the mutant phenotypes, and that these genes function together in a common pathway not previously associated with limb development or embryonic patterning. Conditional knockout of *Wasl* in the mouse demonstrates that this pathway is required for normal limb patterning, and its absence results in a range of skeletal phenotypes that match those seen in *Hoxall* knockout mutants. Through generation of triple loss-of-function *hoxll* zebrafish mutants by CRISPR-Cas9 gene editing, we demonstrate that *waslb* requires *hoxll* function to form supernumerary elements, and more broadly interacts with *hox* genes to pattern the PD fin axis. Our results suggest that zebrafish retain the capacity to form structures with an intermediate regional address specified by *hoxall*, similar to the limb zeugopod. This inherent capacity is activated by simple genetic changes, revealing a novel appendage-patterning pathway that may have played unrecognized roles in the fin to limb transition.

## Results

### Breaking the teleost fin ground plan

Wild-type zebrafish pectoral fins are representative of the ancestral teleost configuration, with four proximal radials arranged side-by-side followed by small nodular distal radials (Fig. 1B). All four proximal radials are in direct contact with the shoulder early in development, but the two posterior radials shift distally in adulthood (*14*). To identify genes affecting the form and pattern of the adult skeleton in zebrafish, we conducted a forward mutagenesis screen focused on dominant mutations (*l3*). In our screen, we isolated a mutant that breaks from the teleost ground plan by forming supernumerary long bones along the PD axis of the pectoral fin endoskeleton (Fig. 1C). We call these new bones ‘intermediate radials.’ They are found between the posterior proximal radials and distal radials, and they do not articulate directly with the shoulder. The overall pattern and size of the fin rays are not dramatically altered. We named this mutant *rephaim (reph)*. The mutant phenotype exhibits varied expressivity and dose sensitivity, as homozygous fish are highly dysmorphic (Figure S2). Due to the severity of the homozygous condition, we have focused our analysis on heterozygous *reph* mutants.

### Uncovering novel fin patterning genes

Using whole-exome sequencing, we mapped the *reph* mutation to chromosome 4 (Figure S3) (15). Analysis of sequence data from the linked interval identified a missense mutation in the gene *wiskott-aldrich syndrome-like b (waslb)*, changing a serine to a proline at amino acid position 265 (Fig. 1D). Waslb is a member of the WAS protein family of actin cytoskeleton signaling regulators, which interact with Cdc42 and Arp2/3 to nucleate filamentous actin formation, and have roles in cell migration, cell signaling, and cell polarity *(16)*. Additionally, nuclear-localized Wasl is required for the transcription of select genes through its interaction with nuclear actin, transcription factors, and RNA polymerase *(17, 18)*.

To verify that the *waslb* mutation causes the *reph* phenotype, as well as to determine the genetic nature of the dominant *reph* allele, we used CRISPR-Cas9 to generate targeted frameshift mutations in wild-type *waslb* as well as in *cis* to the candidate *reph* S265P mutation, and assessed their effect on fin skeleton patterning (Fig. 1D, Figures S4, S5). Loss-of-function of wild-type *waslb* does not have any obvious effect on fin patterning. However, frameshift mutations generated upstream in *cis* to the missense mutation lead to reversion of the mutant phenotype, restoring the wild-type radial pattern. Small, in-frame deletions in *cis* to the mutation failed to rescue the wild-type phenotype. Together, these results demonstrate that the identified S265P mutation in *waslb* causes the *reph* phenotype, likely through a gain-of-function effect. The altered residue in *reph* is conserved across vertebrates and is positioned in a hinge region that is thought to regulate auto-inhibition and localization of the protein *(19)*. Consistent with this, introduction of the *reph* mutation into fluorescently tagged Wasl protein led to a marked reduction in nuclear localization of the fusion protein in cell culture (Figure S5).

### Genetic interactors of Wasl signaling

As *waslb* is a novel factor in appendage development, we sought to identify additional loci that affect fin endoskeleton patterning in a similar manner. Although large dominant screens are not common in zebrafish, mutants having viable dominant phenotypes have been identified as a by-product of recessive genetic screens. We searched published screens and identified a mutant called *wanda (wan)* with a phenotype similar to *reph* (Figure S2) *(12, 20)*. We obtained the *wan* mutant and found that it exhibited intermediate radials in a pattern similar to *reph* (Fig. 1E), a phenotype that had not been previously reported. Similar to *reph, wan* exhibits variable phenotypic expressivity and dose sensitivity (Figure S2).

We mapped *wan* and identified a missense mutation in a conserved residue of the *vav2* gene on chromosome 21 (S67A; Fig. 1D, Figure S3). The Vav family of protooncogenes function as guanosine nucleotide exchange factors for Rho GTPases, and have roles in signal transduction, cytoskeletal regulation, cell motility, and receptor endocytosis *(21-23)*. We created *vav2* loss-of-function mutations using CRISPR-Cas9 and found that *vav2* homozygous null zebrafish are viable and have no skeletal phenotypes (Figures S4, S5), consistent with the phenotype reported for the *Vav2* knockout mouse *(24)*. In addition, truncating mutations made in *cis* upstream of the S67A *wan* mutation cause reversion of the *wan* skeletal phenotype (Fig. 1D, Figure S5). Thus, similar to *reph, wan* causes fin patterning changes through a gain-of-function mutation.

Vav2 activates the small G-protein Cdc42 (25), and Cdc42 in turn regulates the activity of Wasl *(26)*. As such, it is likely that the similar phenotypes observed in *reph* and *wan* mutants are due to their effect on a common signaling pathway. To test this hypothesis, we crossed *waslb* loss-of-function alleles to the *wan* mutant and found that *waslb* function is required for the expression of the *wan* phenotype (Fig. 1E). Thus, *waslb* is epistatic to vav2, and these genes act together in a common pathway to pattern the fin endoskeleton (Fig. 1F). Whole mount *in situ* hybridization reveals that both genes are expressed in fins during early patterning stages (Fig. 1G), suggesting that these genes might act in the developing fin mesenchyme. Interestingly, neither *waslb* nor *vav2* have previously been associated with fin or limb development, skeletogenesis, or embryonic patterning at large.

### Fin fold integrity is unchanged in *rephaim*

In the fin to limb transition, there was a reduction of the dermal skeletal fin rays concomitant with the elaboration of the endoskeleton, until fin rays were ultimately lost in crown tetrapods. Fin rays arise from the larval fin fold (27). Mechanistic hypotheses regarding the fin-to-limb transition posit a developmental trade-off between the fin endoskeleton and the fin fold, such that early formation of the fin fold in teleosts results in their diminutive endoskeleton (28). Previous studies have tested if reduction of the fin fold can result in limb-like characteristics in developing zebrafish fins. Experimental perturbation of the fin fold during early development leads to an enlargement of the endoskeletal disc in larval zebrafish, suggesting that the fin fold represses disc growth *(29, 30)*. However, similar to the effect of reducing *hox13* paralog function (11), subsequent elaboration of the adult fin endoskeleton is generally negligible, or has not been assessed, in the different manipulations. To determine if changes in fin fold dynamics underlie intermediate radial development, we examined fin fold formation in *reph*^+/−^ mutants (Fig. 2). Immunolabeling of actinodin, the collagens that support the larval fin fold, revealed no differences between wild-type and *reph^+/−^* pectoral fins at 3 days post fertilization (dpf; Fig. 2A, B). We also find no difference in endoskeletal disc size at 7 dpf (Fig. 2C-E). The absence of changes in the endoskeletal disc and fin fold is consistent with a lack of patterning defects in 3-week-old mutant fins (Fig. 2F, G). Thus, contrary to previous hypotheses, changes to fin fold development are not required for PD elaboration of the endoskeleton in *reph*^+/−^ mutants.

**Figure 2.**
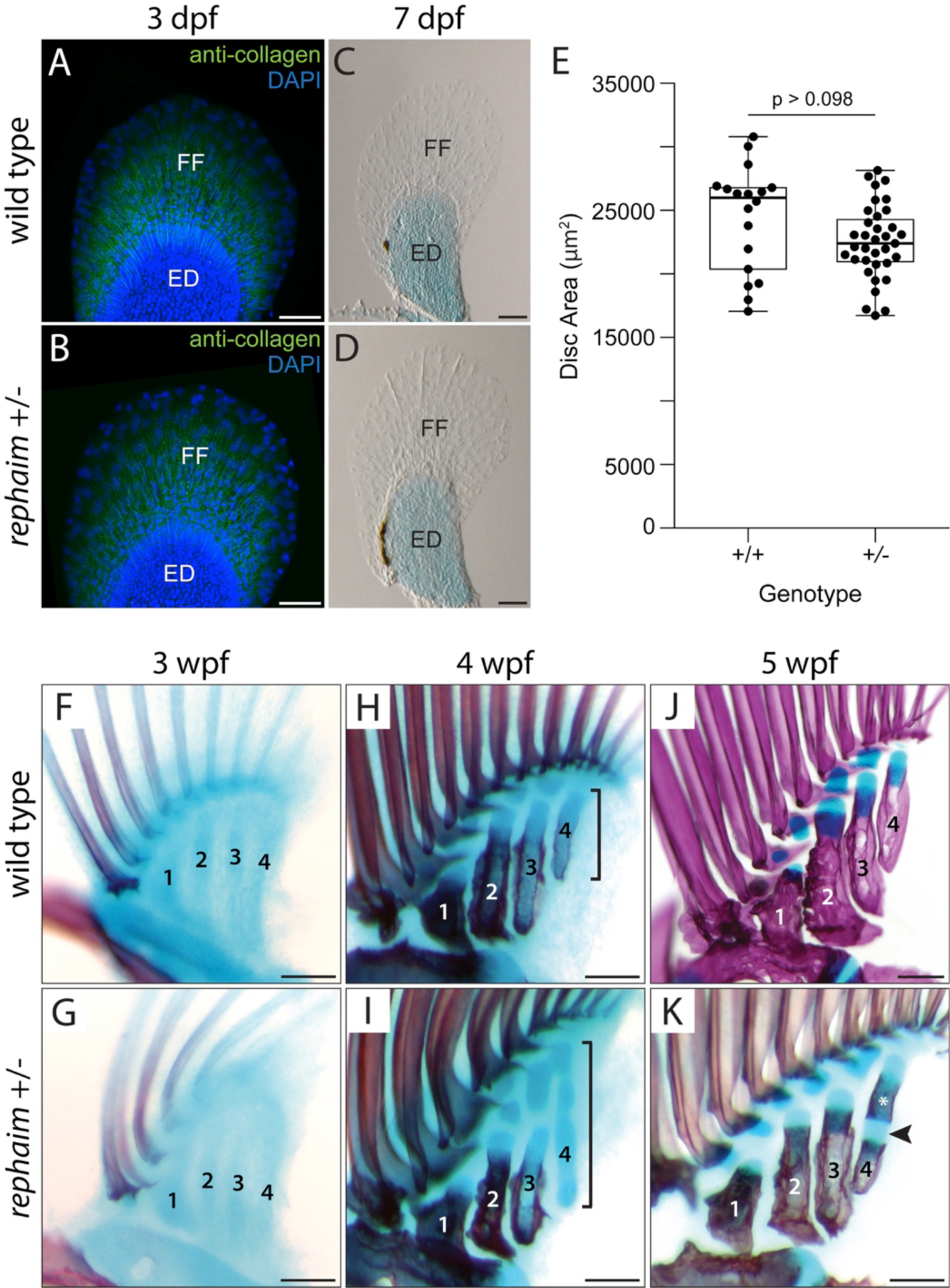
Novel skeletal elements arise by segmentation of a common cartilage precursor. (**A**, **B**) Anti-collagen immunolabeling of actinotrichia fibers (n=4 for both genotypes). (**C**, **D**) Alcian-stained pectoral fins. (**E**) Endoskeletal disc area at 7 dpf is not significantly different between wild-type and *reph*^+/−^ fish (wild type n=18, *reph*^+/−^ n=35, Welch’s Two Sample t-test, *t*=−1.7145, p > 0.098). (**F-K**) Fin skeleton ontogeny in wild-type (**F**, **H**, **J**) and *reph ^+/−^* (**G**, **I**, **K**) fish (n=5 for both genotypes at each age). (**F**, **G**) Separation of the endoskeletal disc into four radial precursors. (**H**, **I**) Proximal mineralization of the radials. Posterior radial cartilages are elongated in *reph^+/−^* mutants (brackets). (**J**) wild-type proximal radials exhibit continuous zones of mineralization. (**K**) *reph^+/−^* mutants develop a secondary bone collar (asterisk) separated from the proximal bone by cartilage (arrowhead). Anterior to left, distal to top; ED, endoskeletal disc; FF, fin fold; 1-4, proximal radials; scale bars (A-D) 50 μm, (F-K) 125 μm.

### Formation of intermediate radials in *rephaim*

In teleost fishes, the fin endoskeleton first forms as a continuous endoskeletal disc of cartilage that is subsequently subdivided into the four proximal radial anlagen by localized involution and trans-differentiation, followed by perichondral ossification (Fig. 2F, H) *(14, 31)*. This process occurs normally in *reph^+/−^* mutants (Fig. 2G). It is only after initial patterning and differentiation that differences are seen between wild-type and mutant fish endoskeletons, when *reph^+/−^* mutants show an elongation of the more posterior cartilage condensations compared to wild-type fish (Fig. 2I). The extended condensations in *reph^+/−^* mutants form multiple sites of mineralization, with the nascent intermediate radial forming a distinct bone collar (Fig. 2K), while wild-type siblings have a continuous zone of mineralization beginning proximally and growing distally (Fig. 2J). The elongated condensation in mutants is subsequently segmented, resolving into two distinct bones separated by joint territories. This mode of ossification and cartilage segmentation in *reph^+/−^* mutants has not been reported in the pectoral fins of any other teleost, and is similar to tetrapod limb development, where an initially continuous condensation is segmented along the PD axis to produce independent elements separated by a joint (32).

### Morphological integration of an expanded fin skeleton

Histological analysis of adult fins reveals that, while proximal radials possess epiphyses only at their distal ends, *reph* intermediate radials form both proximal and distal epiphyses (Fig. 3A, B). Both distal and proximal epiphyses exhibit normal differentiation of the cartilage in *reph^+/−^* mutants. Like the epiphyses of wild-type proximal radials and the long bones of tetrapod limbs, the newly formed epiphyses of intermediate radials express bone patterning genes such as *indian hedgehog a (ihha;* Fig. 3C) (33). Intermediate radials also show evidence of vascular invasion (Fig. 3D), suggesting the activation of endochondral ossification programs in *reph^+/−^* mutants similar to those found in tetrapod long bones and the bones of larger fishes (*34*). In limbs, synovial joints are formed between long bone elements (32). Similarly, in *reph^+/−^* mutants, proximal and intermediate radials form a distinct joint pocket between them with specialized differentiation of mesenchyme shaping the interface (Fig. 3E). We find that the terminal joint marker *prg4b* (35, *36)* is expressed in the superficial chondrocytes of epiphyses, as well as in cells lining the joint space (Fig. 3F). These findings suggest that the joint differentiation programs found in limbs are also activated in the *reph* intermediate radials.

**Figure 3.**
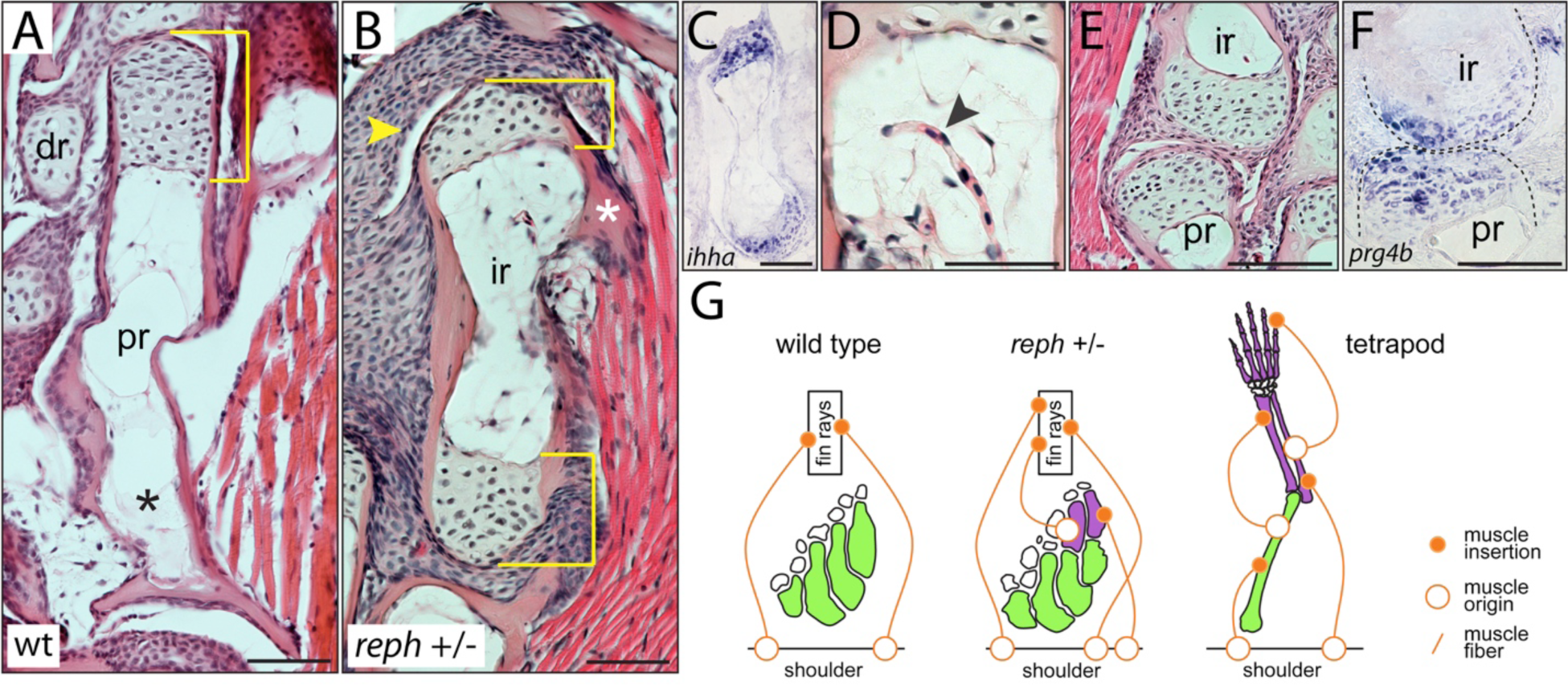
Intermediate radials are long bones morphologically integrated into the fin skeleton. (**A-D**) Fin endoskeleton histology in adult wild-type and *reph^+/−^* adults. (**A**) Wild-type proximal radial with a single distal epiphysis (bracket), and no proximal epiphysis (asterisk). (**B**) Intermediate radial of *reph^+/−^* mutants have dual epiphyses (brackets), a muscle insertion point (asterisk), and a synovial cavity (arrowhead). (**C**) *in situ* hybridization of chondrogenic marker *ihha* in dual epiphysial compartments of an intermediate radial. (**D**) Blood vessel invading an intermediate radial (arrowhead). (**E**) Histology of joint between intermediate and proximal radials. (**F**) *in situ* hybridization of terminal joint marker *prg4b* expression in the proximal radial-intermediate radial joint. (**G**) Diagram of muscle attachment in the tetrapod limb versus the pectoral fins of wild-type and *reph* mutant zebrafish. *Green*, skeletal elements attaching to the shoulder; *purple*, secondary branching elements; *white*, distal radials and carpals; dr, distal radial; ir, intermediate radial; pr, proximal radial; anterior to left, distal to top; scale bars 50 μm.

In limbs, muscles originate from the shoulder as well as the limb bones and insert on more distal positions along the appendage (Fig. 3G). Zebrafish, in contrast, have seven muscles that originate on elements of the shoulder girdle and insert directly on the dermal fin rays, bypassing the fin long bones entirely (37). This configuration of the pectoral fin musculature is representative of most teleost fishes, and only in certain derived lineages does a muscle insert on pectoral fin radial bones (35). Surprisingly, *reph^+/−^* intermediate radials exhibit novel insertion points from muscles originating from the shoulder (Fig. 3B). Thus, not only does the *reph* mutation reveal a capacity to make differentiated and patterned long bones, but these elements are morphologically integrated to form limb-like joints and muscle connections not seen in the fins of other teleosts.

### *Wasl* is required for normal limb patterning and axial identity in tetrapods

Both *wasl* and *vav2* have not previously been implicated in the regulation of developmental patterning. To assess if the patterning roles of *vav2/waslb* signaling are zebrafish specific, or if they are conserved across bony fishes, we asked if *Wasl* is required for normal tetrapod limb development. As homozygous somatic knockout of *Wasl* in mouse is embryonic lethal *(39)*, preventing analysis of adult limb patterning, we generated mice with conditional loss of *Wasl* function in developing limb progenitors by crossing the *Prrx1-Cre* driver *(40)* into a floxed *Wasl ^L2L^* background (41). Wasl^L2L/^+ and Wasl^L2L/L2L^ animals lacking the Cre driver are phenotypically wild type, as are Wasl^L2L/^+ heterozygotes with the driver (Fig. 4A-D, Supplemental Table 1). However, Wasl^L2L L2L^; *Prrx1-Cre* limb knockout (LKO) mice exhibit dramatic defects in limb and axial patterning. The long bones of LKO mouse limbs are shorter and wider than those of wild-type siblings (Fig. 4A, C). LKO mice also exhibit various coalitions of the carpal bones (Fig. 4B), while the tibia and fibula fail to fuse (Fig. 4C). Axial homeosis occurs in LKO mice, where the first sacral vertebra is transformed to exhibit partial or complete lumbar identity (Fig. 4D). This demonstrates that *Wasl* is necessary for proper patterning in limbs and the vertebral column. Intriguingly, this syndrome of axial and appendicular phenotypes closely matches those resulting from somatic loss of *Hoxall* function *(42)*, a gene thought to have played a key role in the fin to limb transition.

**Figure 4.**
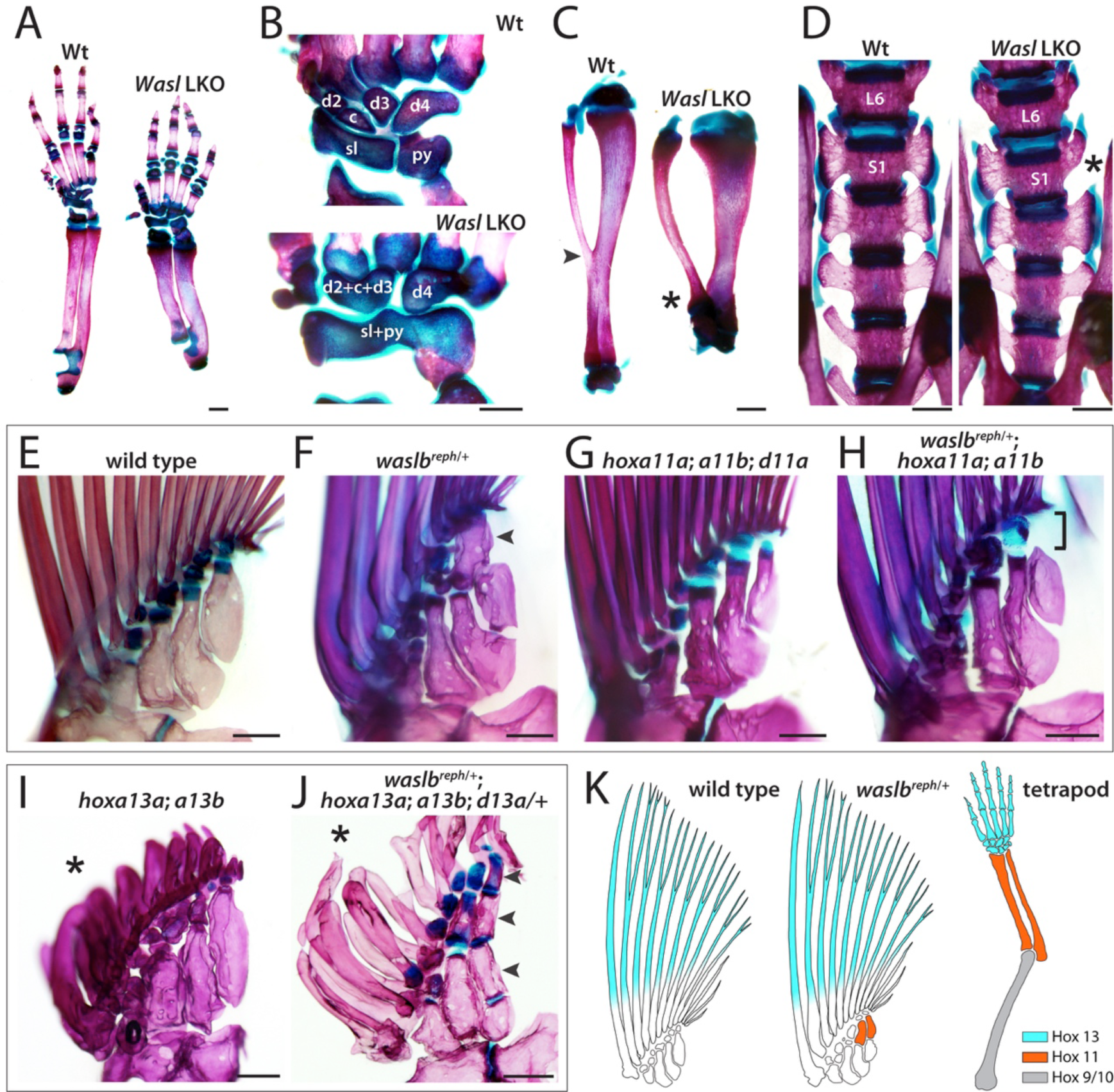
*Wasl* is required for normal limb patterning and interacts genetically with Hox genes to pattern fins. (**A-D**) Skeletal analysis of *Prrxl-Cre*; Wasl^L2L/L2L^ (LKO) mice. (**A**) Limb phenotype of LKO and wild-type littermates. (**B**) Carpal fusions in the wrist and (**C**) reduced length and failed fusion of tibia and fibula in LKO mice. (**D**) Homeosis in sacral vertebrae in LKO mice. (**E-J**) Genetic interaction between *waslb* and *hox* genes in zebrafish fin patterning. (**E**) wild-type adult. (**F**) *waslb reph^+/−^* mutants with wild-type *hoxll* paralogs form intermediate radials (arrowhead; n=9, 89%). (**G**) *hoxalla^−/−^*; *hoxallb^−/−^*; *hoxdlla^−/−^* mutants have normal pectoral fins. (**H**) *waslb reph^+/−^*; *hoxalla^−/−^*; *hoxallb^−/−^* mutants fail to form intermediate radials (bracket; n=8, 75%). (**I**) *hoxal3a^+/−^*; *hoxal3b^−/−^* mutants have defective and reduced fin rays (asterisk) but maintain a wild-type proximal radial pattern. (**J**) *waslbreph^+/−^*; *hoxal3a^−/−^*; *hoxal3b^−/−^*; *hoxdl3a^+/−^* mutants have enhanced PD elaboration with multiple intermediate radials (arrowheads). (**K**) Model of region-specific requirements of Hox paralogs in PD patterning of fin and limb bones as revealed by Hox null mutations in zebrafish and mouse. Anterior to left, distal to top; c, central carpal; d2-4, distal carpals; py, pyramidal; L6, lumbar vertebrae 6; S1, sacral vertebrae 1; sl, scapholunate; scale bars (A, C, E) 1 mm, (B) 500 μm, (E-J) 250 μm.

### *Waslb* regulates appendage patterning through *hoxa* function

It is unclear if intermediate radials have a specific positional identity defined by *hox* gene expression, as is observed in the elements of tetrapod limbs. Limbs have a tripartite bauplan that has been canalized through their evolutionary history (Fig. 1), consisting of one long bone (humerus) in the upper arm (the stylopod), two side-by-side long bones (radius and ulna) in the forearm (the zeugopod), and many nodular and long bones in the wrist and hand (the autopod). Differential expression of *Hox* genes along the growing limb is essential for specification of positional identity and growth (9). The A and D members of Hox paralogy group 13 are expressed in the autopod, while the A and D members of group 11 are expressed in the zeugopod *(43-45)*. Loss of either group results in the reduced growth or even absence of the elements in which they are expressed *(46, 47)*. The pectoral fins of teleosts and other ray-finned fishes do not exhibit comparable *hox* regulation, as *hoxll* and *hox13* paralogs are expressed in overlapping domains *(48-50)*. This has been thought to be consistent with the simple adult skeleton formed in teleost fins (48). Interestingly, *hox13* genes are however required for the development of the distal fin rays (*11*).

We asked if intermediate radials formed through activation of *waslb* signaling bear a positional identity specified by a Hox code similar to that found in limbs. Based on the *Wasl* LKO mouse phenotype, we hypothesized that *reph* might interact with *hoxa11* paralogs. To test this, we created null alleles of the *hoxa11a, hoxa11b*, and *hoxd11a* genes and examined how loss of these factors affects the expression of the *reph* phenotype. Homozygous null mutants for all three paralogs are viable and fertile, singularly and in combination, and do not show obvious phenotypes in the formation of the fin skeleton (Fig. 4G, Figure S6). However, in *reph^+/−^* mutants, loss of *hoxa11* paralog function resulted in the loss of intermediate radials, reverting the *reph^+/−^* pectoral skeleton to the wild-type pattern (Fig. 4H). This result suggests that the intermediate radials formed in *reph^+/−^* mutants share a zeugopodial-type identity, and developmental homology, with the radius and ulna. Our findings demonstrate that, in contrast to the roles of *hox13* paralogs in regulating formation of distal fin ray structures (Fig. 4I) (11), *hoxll* paralogs do not have essential roles in normal patterning of the zebrafish fin. Rather, the functional necessity of *hoxall* paralogs is revealed only in the context of intermediate radial formation in *reph^+/−^* mutants.

The *Hox13* group genes can repress the activity of more proximal *Hox* genes in a process termed posterior prevalence (51). In limbs, *Hoxa13* suppresses *Hoxall* activity, and these genes form mutually exclusive expression domains at the stylopod/autopod boundary (52). Consistent with the broad retention of genomic architecture of the Hox complex across vertebrate evolution (53) and the requirement of *hoxll* function to form reph-mediated supernumerary elements, we found that loss of *hox13* function resulted in the development of additional supernumerary long bones, but only in the context of *waslb* gain-of-function caused by the *reph* mutation (Fig. 4J).

The necessity of *hoxall* paralogs for the expression of the *reph^+/−^* phenotype reveals the presence of a functional Hox code and regulatory network (53) within developing teleost pectoral fins capable of specifying intermediate domains similar to that found in limbs. This suggests that the intermediate radials have a *hoxll* identity and share developmental homology with the zeugopodial structures of the tetrapod limb (Fig. 4K). However, the manifestation of this positional code is not normally expressed in teleost fishes, but is revealed by the action of the *reph* and *wan* mutations.

## Discussion

The integration and differentiation of the novel long bones formed in *reph* and *wan* pectoral fins is unique amongst teleost fishes, and harkens to the development and anatomy of long bones in the limbs of tetrapods. The common ancestor of all bony fishes, including humans, had a pectoral fin skeleton that consisted of multiple long bones arranged side by side, each articulating with the shoulder (Figure S1) *(7, 8)*. This polybasal condition of the endoskeleton consisted of unsegmented elements in the middle and anterior portions of the appendage, and the posterior metapterygium, which branched and was segmented along the PD axis. It is hypothesized that the tetrapod limb skeleton evolved via loss of the anterior components and elaboration of the metapterygium. Teleosts, on the other hand, are thought to have maintained the polybasal condition and lost the metapterygium *(54-56)*. Interestingly, the skeletal elaboration seen in *reph* and *wan* is restricted to the posterior region of the pectoral skeleton, resembling the morphology of basal actinopterygian and lobe-finned fishes *(1, 57)*. Thus, it is possible that teleosts retain a cryptic ancestral axis of growth, corresponding to the metapterygial axis, whose potential for elaboration is revealed by these mutants.

It is unlikely that the mutations identified in this study are specifically instructive of limb-ness, but are instead permissive of endogenous limb-like developmental programs latent in fishes. A component of these mutation-activated programs is a functional Hox code, retained in teleost fishes, with specific intermediate *(hoxall)* and distal *(hoxa13)* patterning cues. Our results reveal latent or emergent properties of development within vertebrate appendages to form elaborate, articulated skeletal structures. These patterning processes were present in the bony fish ancestor and potentially refined in the evolution of lobe-finned fishes during the transition to land.

## Methods

### Zebrafish husbandry, strains, and mutagenesis

All zebrafish lines were maintained and propagated as described by Nuesslein-Volhard & Dahm (55). This study was conducted with ethical approval from the Institutional Animal Care and Use Committee of Boston Children’s Hospital. A complete description of the husbandry and environmental conditions for the fish used in these experiments is available as a collection in *protocols.iodx.doi.org10.17504protocols.io.mric54n. rephaim (dmh22)* was isolated as part of a broad mutagenesis screen performed at Boston Children’s Hospital *(13). wanda (ty127)* was initially isolated and described in the original Tuebingen screens (12). *wanda* mutant stocks were obtained from the European Zebrafish Resource Center (http://www.ezrc.kit.edu/).

### Mapping rephaim and wanda mutant zebrafish

Whole-exome sequencing was used to map the loci and mutations causing the *reph* and *wan* phenotypes through homozygosity-by-descent methods (15). As both mutants exhibit dose-dependent phenotypes, homozygotes could be isolated and used for mapping. For each mutant, genomic DNA from 25 F2 homozygous mutants stemming from a WIK outcross was extracted from whole larvae using the Qiagen DNeasy Blood & Tissue kit. 3 μg of pooled DNA was sheared to an average fragment size of 200 bp using a Covaris E220 Focused Ultrasonicator. Barcoded sequencing libraries were prepared from fragmented DNA using the KAPA Hyper Prep Kit following the manufacturer’s protocol. We used Agilent SureSelect DNA target enrichment baits designed using the Zv9 genome assembly to enrich for coding regions of the zebrafish genome (59). 50 bp single-end sequencing was performed on an Illumina HiSeq machine, resulting in average exome coverage of 7x for *rephaim* and 13x for *wanda*. Fine mapping of linked intervals was performed with polymerase chain reaction (PCR)-based analysis of recombinants using unique single nucleotide polymorphisms (SNP) and simple sequence length polymorphisms (SSLP). Mapping PCR primers are listed in Supplemental Table 2.

### Gene editing and isolation of frameshift mutations

CRISPR-Cas9 site-directed mutagenesis was used to generate null alleles for each target gene. Suitable target sites located early in the first exon of each gene were selected using CHOPCHOP (http://chopchop.cbu.uib.no/).

hoxa11a_1: 5’ACTGGCACATTGTTATCCGT 3’; hoxa11b_1: 5’ CGTCTTCTTGCCCCATGACA 3’;

hoxa11b_2: 5’ TTTGATGAGCGGGTACCCGT 3’; hoxd11a_1: 5’ CGCTTCGTACTATTCAACGG 3’;

hoxd11a_2: 5’ CTATTCTTCGAACATAGCGC 3’; vav2_1: 5’ CTGAGGGCCAAACGACCCGG 3’;

vav2_2: 5’ TGGAGGAGTGGAGGCAGTGC 3’; waslb_1: 5’ ATAGAGCCAACATTGAGCGC 3’;

waslb_2: 5’ AGAGCACTTCGTTTTCCTGA 3’; waslb_3: 5’ GCGGTCCACCTTCAAACAGG 3’.

Guides were synthesized by IDT (Integrated DNA Technologies, Coralville, IA) for use with their Alt-R CRISPR-Cas9 system. Gene-specific guides were multiplexed and injected at a concentration of 6.25 μM each crRNA guide and 500 ng/μL Cas9 mRNA into single-cell zebrafish embryos. Progeny of injected fish were screened for the presence of inherited lesions resulting in frameshifts and truncations, and these progeny were used as founders of stable mutant lines. Genotyping primers and strategies for each allele are reported in Supplemental Table 2. Amino acid sequence alterations caused by each lesion are reported in Supplemental Table 3.

### Skeletal staining and histology

Adult fish were fixed overnight at room temperature with agitation in 3.7% formaldehyde in phosphate buffered saline (PBS) pH 7.4. After fixation, fish were rinsed briefly with PBS and stepped through one-hour ethanol/distilled water washes, with 30%, 50%, 70%, 95%, and finally 100% ethanol. Cartilage was then stained overnight at room temperature with 0.015% Alcian Blue GX in 30% acetic acid in ethanol. One-hour ethanol/distilled water washes were then used to move the animals through 100%, 70%, 50%, 30% ethanol, and then washed twice in tap water. Fish were then macerated using 0.15% trypsin in 65% saturated sodium borax solution at 37°C. Typically fish were digested for 1 to 3 hours, stopping the process once the caudal peduncle and anal fin radials were visible when held up to light. Mineralized tissues were stained with 0.25% Alizarin Red S in 0.5% KOH (potassium hydroxide) overnight. If one night of staining was inadequate, a second overnight wash was performed with fresh staining solution. For storage and imaging, stained fish were then moved through a glycerol/0.5% KOH series using overnight washes of 20%, 40%, 60%, 80% glycerol in 0.5% KOH, and finally 100% glycerol. Larval and juvenile fish were stained with the same protocol, with the exception that larval fish were left in the cartilage stain for only 4 hours. All wash volumes are 20 mL and all steps were performed in scintillation vials.

### Calculations of endoskeletal disc area

Wild-type and *reph^+/−^* 7 dpf larvae were stained with Alcian blue. After staining, pectoral fins were removed and the endoskeletal disc was measured using integrated measurement tools in the Nikon Imaging Software. For each fin, the disc area was measured in triplicate and the average taken to be used in statistical analysis. Statistical comparison of average disc size was performed by Welch’s Two Sample t-test in the R statistical module *(60)*.

### Whole-mount actinotrichia collagen labeling

Whole-mount immunolabeling of actinodin collagen filaments and DAPI counter-labeling was carried out as previously described by Lalonde and Akimenko (2018) *(29, 30)*.

### Histology

Fish were fixed overnight in 3.7% formaldehyde in PBS at room temperature. After fixation, the pectoral girdle was removed and decalcified in 10% EDTA (ethylenediaminetetraacetic acid) for several days, with daily solution changes. Following decalcification, fins were embedded in paraffin using standard practices. Hematoxylin and Eosin staining was performed using standard protocols.

### *In situ* hybridization

Whole mount *in situ* hybridization was performed following Jackman *et al. (61)* with minor modifications. Embryos were pre-treated with 2.5 μg/mL proteinase K for 30 minutes at room temperature. *In situ* hybridization on sections was performed following Smith *et al*. (62). Prior to embedding in OCT (Optimal Cutting Temperature, Sakura), adult fins were decalcified overnight in 10% EDTA pH 7.6.

### Cell localization constructs and assay

The gene fusion construct mEmerald-N-Wasp-C-18, consisting of an N-terminal GFP fused to the *Mus musculus Wasl* coding region driven by the CMV promoter, was a gift from Michael Davidson (Addgene plasmid #54199). Site-directed mutagenesis via the QuikChange II XL kit (Agilent) was used to introduce the S249P mutation with the primers 5’CCCAGCTTAAAGACAGAGAAACCTCAAAAGTTATTTATGACTTTATTG’3 and 5’CAATAAAGTCATAAATAACTTTTGGTGTTTCTCTGTCTTTAAGCTGGG’3. HeLa cells were seeded on 6-well plates in Dulbecco’s Modified Eagle’s medium (DMEM, Life Technologies) supplemented with 10% fetal bovine serum (FBS, Life Technologies) and 1000 U penicillin/streptomycin (Life Technologies). Cells were transfected using FuGENE transfection reagent (Promega) with either 1 μg wild-type or S249P construct. Two days after transfection, cells were given fresh media, stained with NucBlue™ Live Cell Stain (Molecular Probes), and photographed using a Zeiss EVOS imaging system.

### Mouse husbandry and breeding

Mice were maintained in the animal resources facilities at Boston Children’s Hospital. This work was conducted with the ethical approval of the Institutional Animal Care and Use Committee at Boston Children’s Hospital. Adult mice homozygous for the conditional *Wasl* allele *(Wasl^L2L^)* were a gift from Dr. Scott Snapper. These mice were crossed to the *Prrxl-Cre* driver line. All resulting progeny were phenotypically wild type, and those carrying the driver transgene were both in-crossed to one another and crossed to Wasl^L2L/L2L^ homozygotes. Offspring genotypes were observed in the expected ratios and were adult viable. Only Wasl^L2L/L2L^; *Prrxl-Cre* animals expressed an abnormal phenotype.

## Acknowledgements

The authors would like to thank Drs. Joana Lopes and Matthew Warman for assistance in cell culture and mouse husbandry, Dr. Neil Shubin for providing *hox13* zebrafish mutants, and Dr. Scott Snapper for sharing the *Wasl* conditional mouse. The authors would like to thank Drs. Randall Dahn, Seth Donoughe, and James Hanken for their comments on early drafts of the manuscript. This work was supported in part through support from Children’s Orthopedic Research Foundation, graduate research funds from the department of Organismic & Evolutionary Biology at Harvard University to MBH, and NSF DDIG 1600920.

## Author Contributions

MBH, KH, and MPH generated materials, designed the genetic and experimental approaches, interpreted results, and wrote the paper.

## Supplemental Data

**Supplemental Figure 1.**
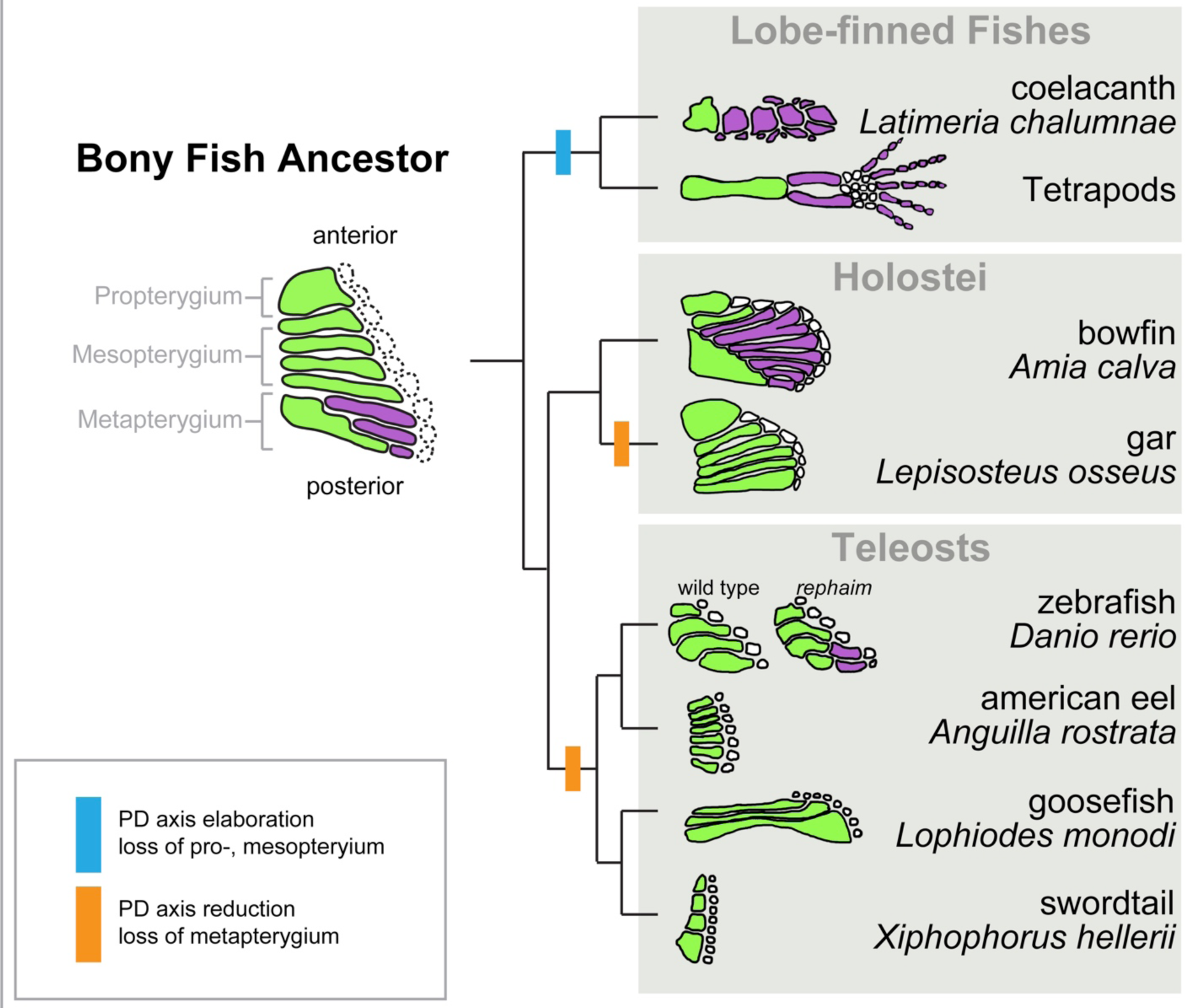
Phylogenetic model of pectoral appendage articulation in bony fishes. The last common ancestor of the bony fishes, the clade that encompasses the teleost and tetrapod lineages, had a polybasal condition in which the most proximal endochondral elements of the fin articulated with the shoulder girdle. The pectoral fin consisted of the anterior propterygium, the middle mesopterygium, and the posterior metapterygium. In the lineage leading to tetrapods, the propterygium and mesopterygium are postulated to have been lost along with the dermal fin rays, while new long bones that articulate end on end were added along the PD axis of the metapterygium, ultimately forming the limb. Conversely, in the teleost lineage, it is proposed that the metapterygium was lost while the propterygium and mesopterygium were retained, and no teleost exhibits end-on-end articulation of long bones in the pectoral fin. While rare, variation in the teleost pectoral fin endoskeleton is observed in proximal radial length as well as the number of radials along the AP axis of the fin in different clades. However, no teleost species showing PD long bone articulation comparable to that seen in tetrapods have been described.

**Supplemental Figure 2.**
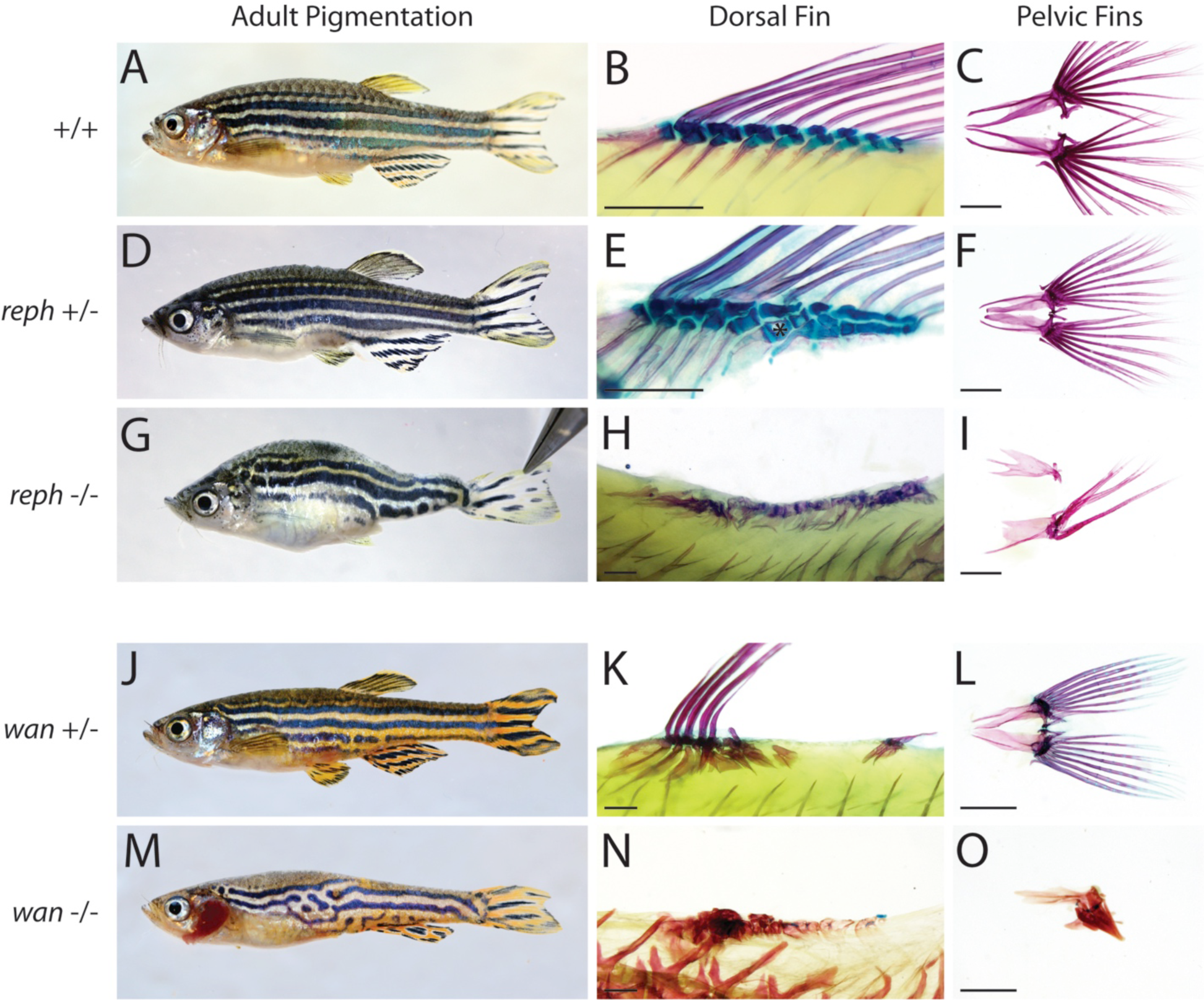
Additional adult phenotypes of *reph* and *wan* mutants. Adult pigmentation and skeletal patterning phenotypes of dorsal and pelvic fins in *reph* and *wan* mutants. (**A-C**) wild-type adult. (**D-F**) *reph* heterozygotes show increased number of radials in the dorsal fin (asterisk), however pelvic fins (**F**) are unaffected. (**G-I**) *reph* homozygous mutants show more severe effect on pigmentation (**G**), and defects in lepidotrichia growth and maintenance (**H**, **I**). Dorsal fin endoskeleton in homozygous fish is uninterpretable. (**J-L**) *wan* heterozygotes show alteration in pigmentation (**J**), and medial fin growth and patterning (**K**). Pelvic fins are unaffected (**L**). (**M-O**) Similar to *reph*, *wan* homozygous fish show a more severe effect on pigmentation (**M**), and highly dysmorphic fin rays and endoskeletal elements in the dorsal and pelvic fins (**N**, **O**). Scale bars (B, E, H, K, N) 500 μm, (C, F, I, L, O) 1 mm.

**Supplemental Figure 3.**
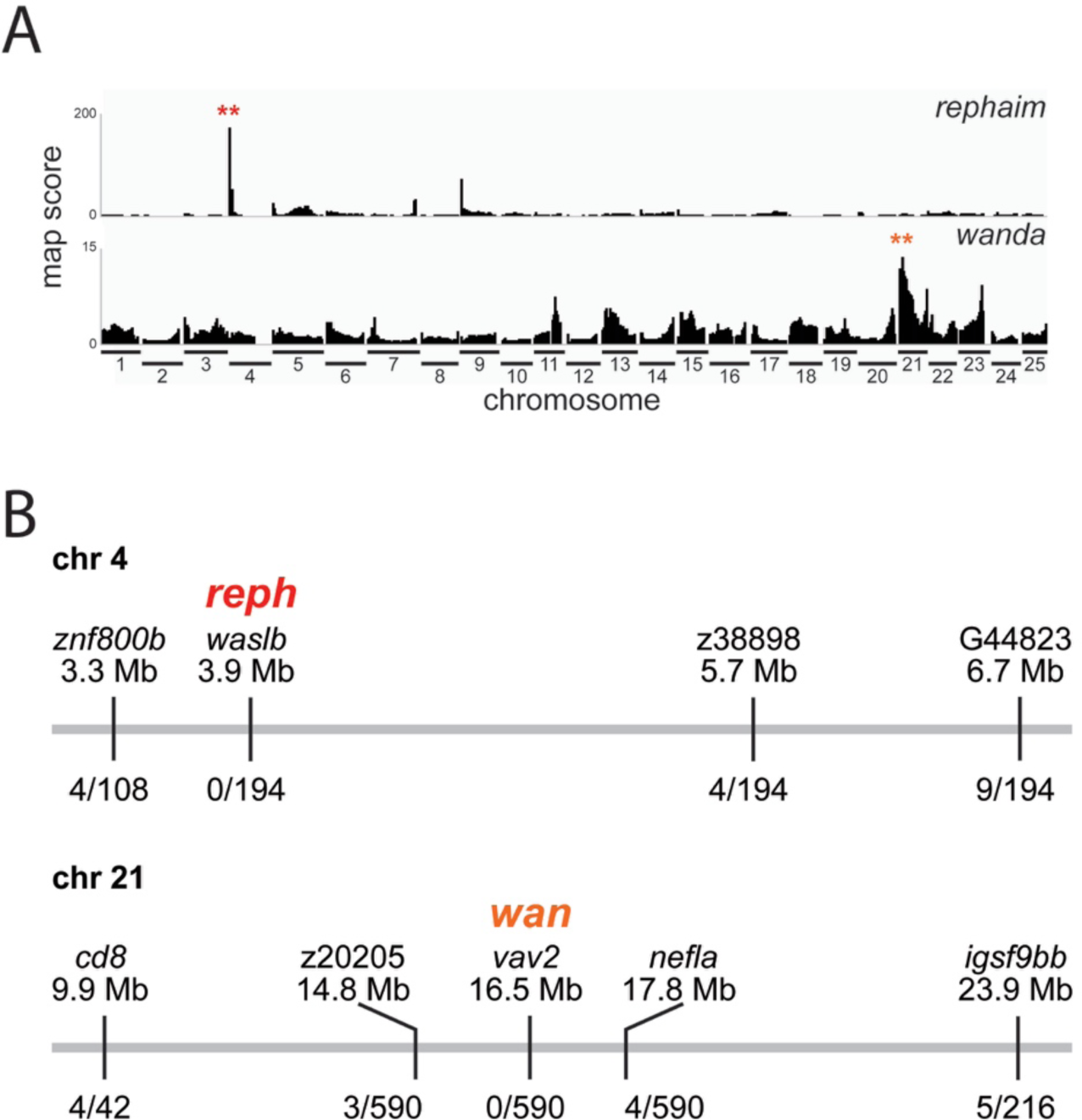
Mapping the causative mutations underlying the *rephaim* and *wanda* >phenotypes. (**A**) Mapping score from homozygosity-by-descent analysis for *reph* and *wan* mutants based on whole-exome sequencing of homozygous populations. Asterisks highlight regions of putative linkage. (**B**) Fine mapping of the chromosome with the highest mapping score by recombination analysis in individual homozygous mutants. Shown are informative markers used, their position, and the number of recombinants identified per meiosis scored. Within the smaller linked interval, sequence data from the mutant pool was used to identify nonsynonymous SNPs as candidate mutations responsible for the altered skeletal phenotype of *reph* and wan.

**Supplemental Figure 4.**
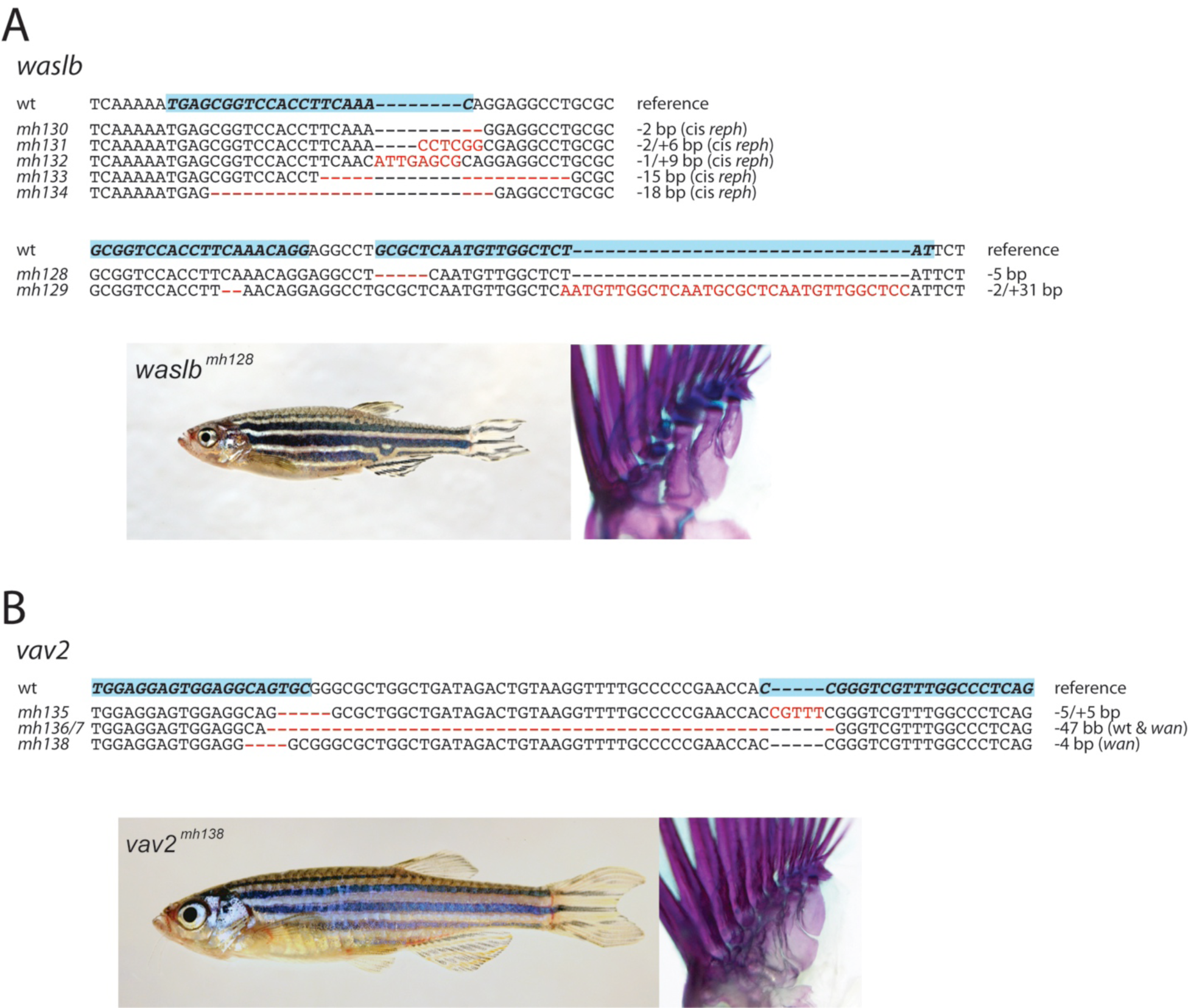
CRISPR-generated loss-of-function mutants of *waslb and vav2*. Mutant lines of (**A**) *waslb* and (**B**) *vav2* generated by CRISPR-Cas9 targeted gene editing. Shown is the targeted wild-type sequence as well as the sequence for the generated mutant alleles (blue indicates guide target sequence). Pictures show representative homozygous mutants. All mutants are homozygous viable and fertile. Whole mount skeletal stained adult pectoral fins from homozygous mutants show no obvious defects in fin patterning.

**Supplemental Figure 5.**
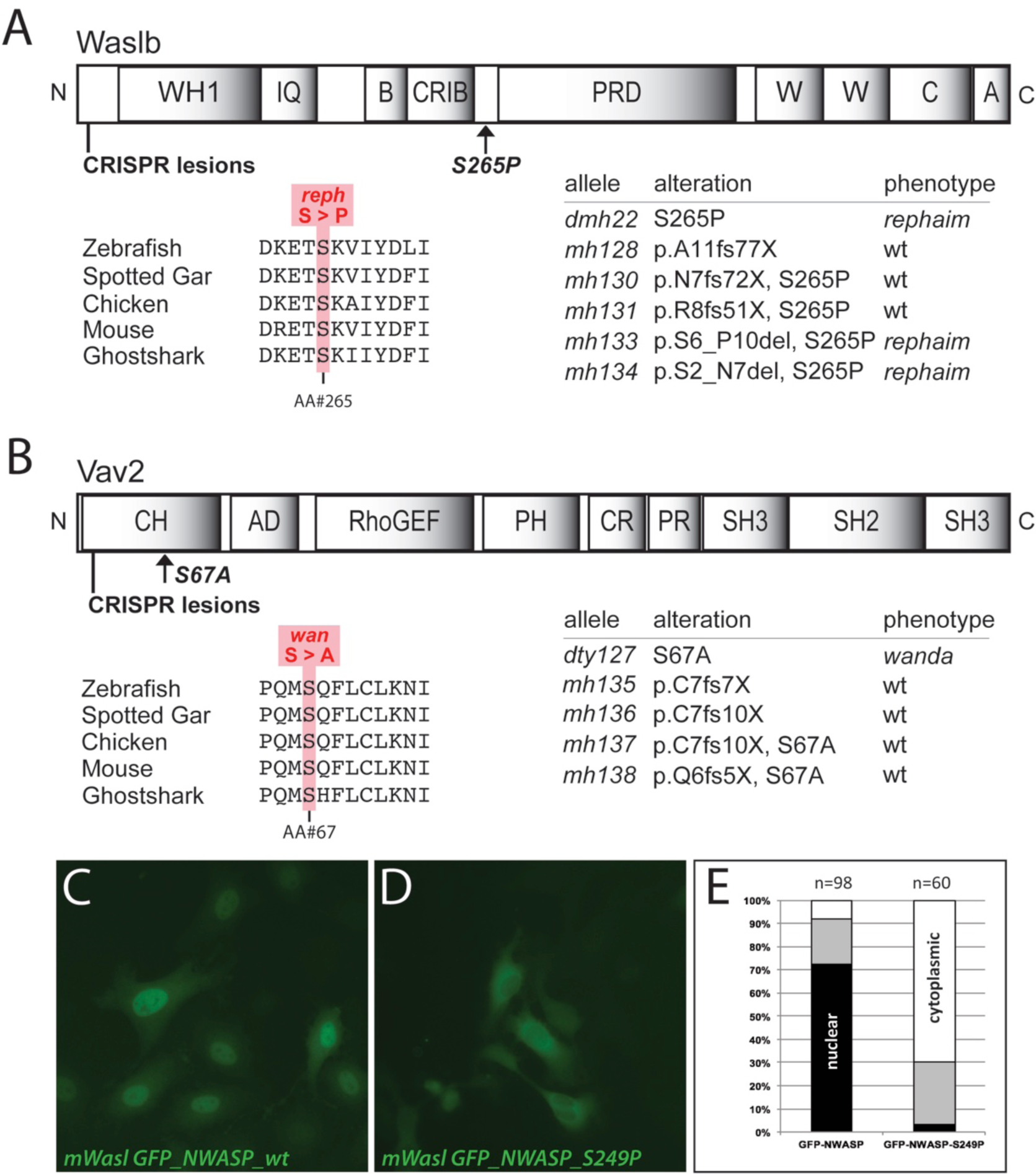
Gain-of-function mutations in *waslb* and *vav2* cause the *reph* and *wan* phenotypes, respectively. (**A**) Waslb and (**B**) Vav2 protein schematics showing positions of missense mutations and CRISPR-generated lesions. Multiple sequence alignment reveals conservation of the mutated residues across vertebrates. Mutant alleles are listed with their consequence on amino acid sequence and their phenotypic effect. (**C-D**) HeLa cells transfected with constructs encoding N-terminal GFP fusions to mouse wild-type N-Wasp (**C**) and an N-Wasp version containing the orthologous *reph* mutation (S249P) (**D**). (**E**) Qualitative scoring of transfected cells; *white*, predominantly cytoplasmic localization; *black*, predominantly nuclear localization; *grey*, no bias observed in localization.

**Supplemental Figure 6.**
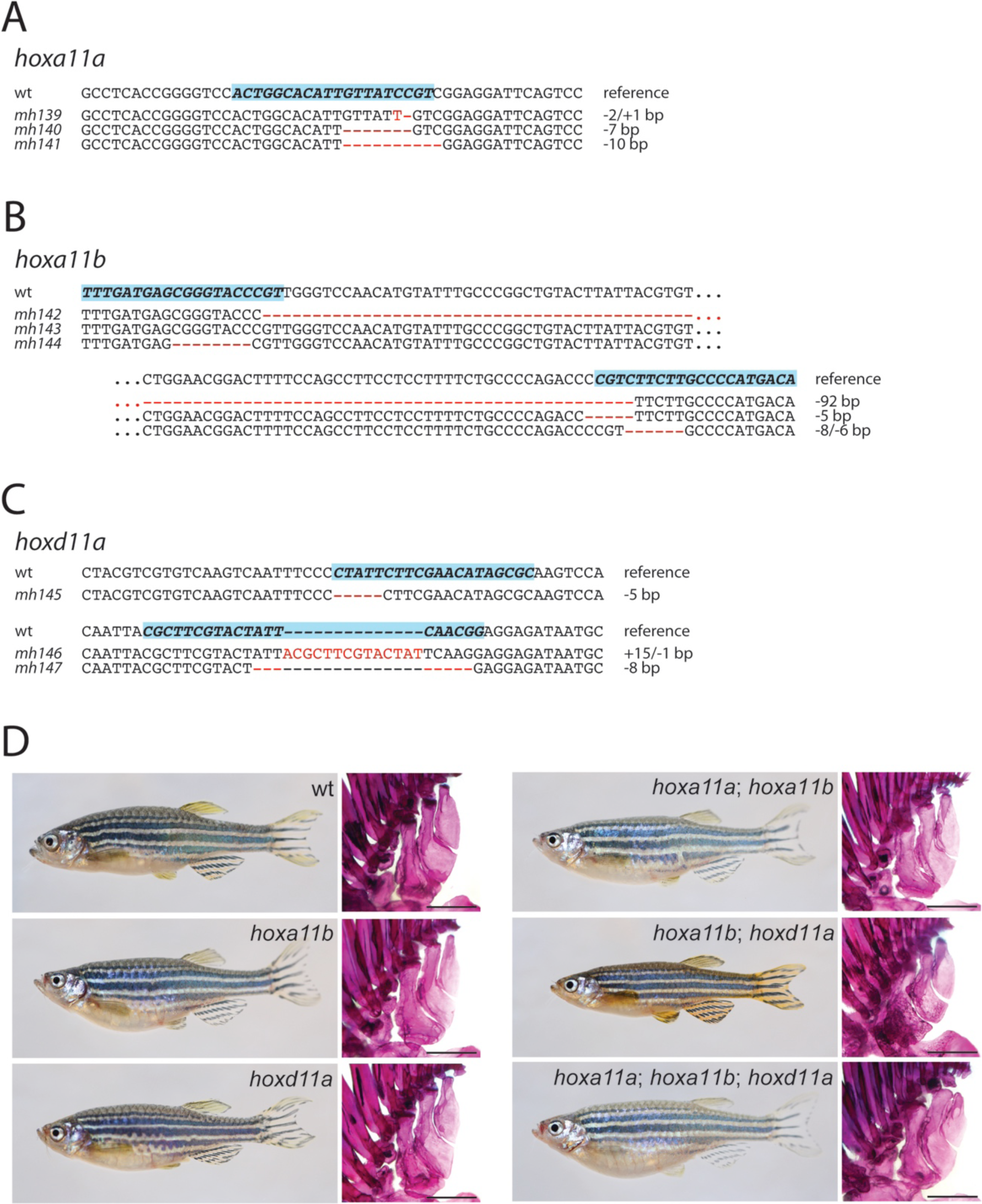
CRISPR-generated *hoxll* loss-of-function mutants. Mutant lines of *hoxll* paralogs (**A**) *hoxalla*, (**B**) *hoxallb*, and (**C**) *hoxdlla* generated by CRISPR-Cas9 targeted gene editing. Shown is the targeted wild-type sequence as well as the sequence for the generated mutant alleles (blue indicates guide target sequence). (**D**) Representative singular and compound mutants. All mutant combinations are viable and fertile. Scale bars 500 μm.

**Supplemental Table 1.** Skeletal phenotypes in *Wasl* LKO mice. Counts reported for limb and axial phenotypes observed in *Wasl* LKO mice and wild-type littermates cleared and stained at P30. c, central carpal; d2, distal carpal 2; d4, distal carpal 4; pi, pisiform; py, pyramidal; sl, scapholunate; T, triangular.

|  | wild type<br>(n=5) | *Wasl* LKO<br>(n=5) |
| --- | --- | --- |
| Vertebral pattern |  |  |
| 6 Lumbar / 4 Sacral (wild type) | 5 | 2 |
| 6 Lumbar / 1 Mixed Identity / 3 Sacral | 0 | 2 |
| 7 Lumbar / 3 Sacral | 0 | 1 |
| Wrist pattern |  |  |
| normal | 5 | 0 |
| d2 + c + d4 fusion | 0 | 5 |
| sl + py fusion | 0 | 1 |
| pi + T fusion | 0 | 5 |
| Tibia and Fibula: normal/unfused | 5/0 | 0/5 |
| Limb long bones: normal/shortened | 5/0 | 0/5 |

